# GM-CSF drives immune-mediated glomerular disease by licensing monocyte-derived cells to produce MMP12

**DOI:** 10.1101/2022.06.13.495915

**Authors:** Hans-Joachim Paust, Ning Song, Donatella De Feo, Nariaki Asada, Selma Tuzlak, Yu Zhao, Jan-Hendrik Riedel, Malte Hellmig, Amirrtavarshni Sivayoganathan, Anett Peters, Anna Kaffke, Alina Borchers, Ulrich O. Wenzel, Oliver M. Steinmetz, Gisa Tiegs, Elisabeth Meister, Maja T. Lindenmeyer, Elion Hoxha, Rolf A.K. Stahl, Tobias B. Huber, Stefan Bonn, Catherine Meyer-Schwesinger, Thorsten Wiech, Jan-Eric Turner, Burkhard Becher, Christian F. Krebs, Ulf Panzer

## Abstract

Glomerulonephritis is a group of immune-mediated diseases that cause inflammation within the glomerulus and adjacent compartments of the kidney and is a major cause of end-stage renal disease. T cells are among the main drivers of glomerulonephritis. However, the T cell subsets, cytokine networks, and downstream effector mechanisms that lead to renal tissue injury are largely unknown, which has hindered the development of targeted therapies.

Here we identify a population of GM-CSF-producing T cells that accumulates in the kidneys of patients with ANCA-associated glomerulonephritis, infiltrates the renal tissue in a mouse model of glomerulonephritis, and promotes tissue destruction and loss of renal function. Mechanistically, we show that GM-CSF producing T cells licence monocyte-derived cells to produce matrix metalloproteinase 12 (MMP12), which cleaves components of the glomerular basement membrane and exacerbates renal pathology. These findings provide a mechanistic rationale for the immunopathology of T cell-mediated diseases and identify the “GM-CSF – monocyte-derived cells – MMP12” pathway as a promising therapeutic target in treatment of glomerulonephritis.

## INTRODUCTION

Glomerulonephritis (GN) is a group of immune-mediated diseases and a major cause of end-stage renal disease (*1*). Glomerular inflammation and injury can be a consequence of systemic autoimmune diseases, such as anti-neutrophil cytoplasmic antibody (ANCA)-associated vasculitis and systemic lupus erythematosus, or kidney-restricted diseases, such as IgA nephropathy (*1–3*). In addition to the role of glomerular autoantibodies and immune deposit formation to the immunopathology, several studies demonstrated the causal role of proinflammatory T cells in glomerulonephritis (GN) (*2, 4–6*). In particular, CD4^+^ T cells orchestrate the immune response by instructing other (immune) cells in the inflamed tissue via secretion of various cytokines (*7*). Based on their different polarization states, corresponding cytokine profiles and primary target cells, three major categories of T helper cell responses can be categorized. The type 1 response is characterized by GM-CSF- and IFN-γ-producing T cells activating and licensing myeloid cells. T cells providing help to B cells, eosinophils, and mast cells for example via IL-4 and IL-13 expression are the key feature of the type 2 response, whereas in the type 3 response IL-17A and IL-22 expressing T cells mainly act on epithelial or stromal cells (*8, 9*).

In the majority of immune-mediated diseases the dominant type of T cell profile, the underlying cytokine signaling pathways and ensuing inflammatory cascades are not yet fully understood. From a clinical perspective, the lack of specific knowledge about the immunopathological role of these immune responses hinders the implementation of pathogenesis-based therapies. This is especially true for patients with crescentic GN, the most aggressive form of autoimmune kidney disease with the worst prognosis (*3*).

New single cell technologies allow high-dimensional immune cell characterization directly from the tissue (*10, 11*) and may allow to more specifically address their function in GN. To decipher the cellular immune response in GN, we investigated the profile of renal T cells and defined a population of GM-CSF-producing inflammatory T cells in patients with ANCA-associated GN (ANCA-GN, the most common cause of crescentic GN) as well as in a crescentic GN mouse model. Our studies reveal that GM-CSF-producing T cells were localized in direct proximity to myeloid cells in the kidney and activated monocyte-derived cells (MdC) to produce matrix metalloproteinase 12 (MMP12), resulting in the degradation of structural elements of the kidney, such as the glomerular basement membrane, and loss of renal function. Molecular and histopathological analyses of kidney biopsies from patients with ANCA-GN displayed the enrichment of GM-CSF inflammatory pathways and the presence of MMP12-expressing MdCs in the inflamed kidney, which correlated with disease activity and renal pathology, highlighting its potential contribution to the pathogenesis of GN.

## RESULTS

### Study cohort and experimental overview

We included three independent cohorts of patients with glomerulonephritis for molecular profiling, pathological examination and clinical analysis in our study (Figure 1A, Supplemental Table 1). First, the Hamburg GN Registry (cohort 1) (*12*), was used as a discovery cohort to analyze CD3^+^ T cells by FACS, single cell RNA (scRNA) sequencing and multiplex staining to define the T cell response in the kidney. Second, we used the European Renal cDNA Bank (cohort 2) (*13*) for molecular profiling across different forms of glomerulonephritis and for the correlation of molecular markers with disease activity. Finally, we used the Clinical Research Unit 228 (CRU 228) ANCA cohort (cohort 3) (*14*) for validation analysis.

**Figure 1:**
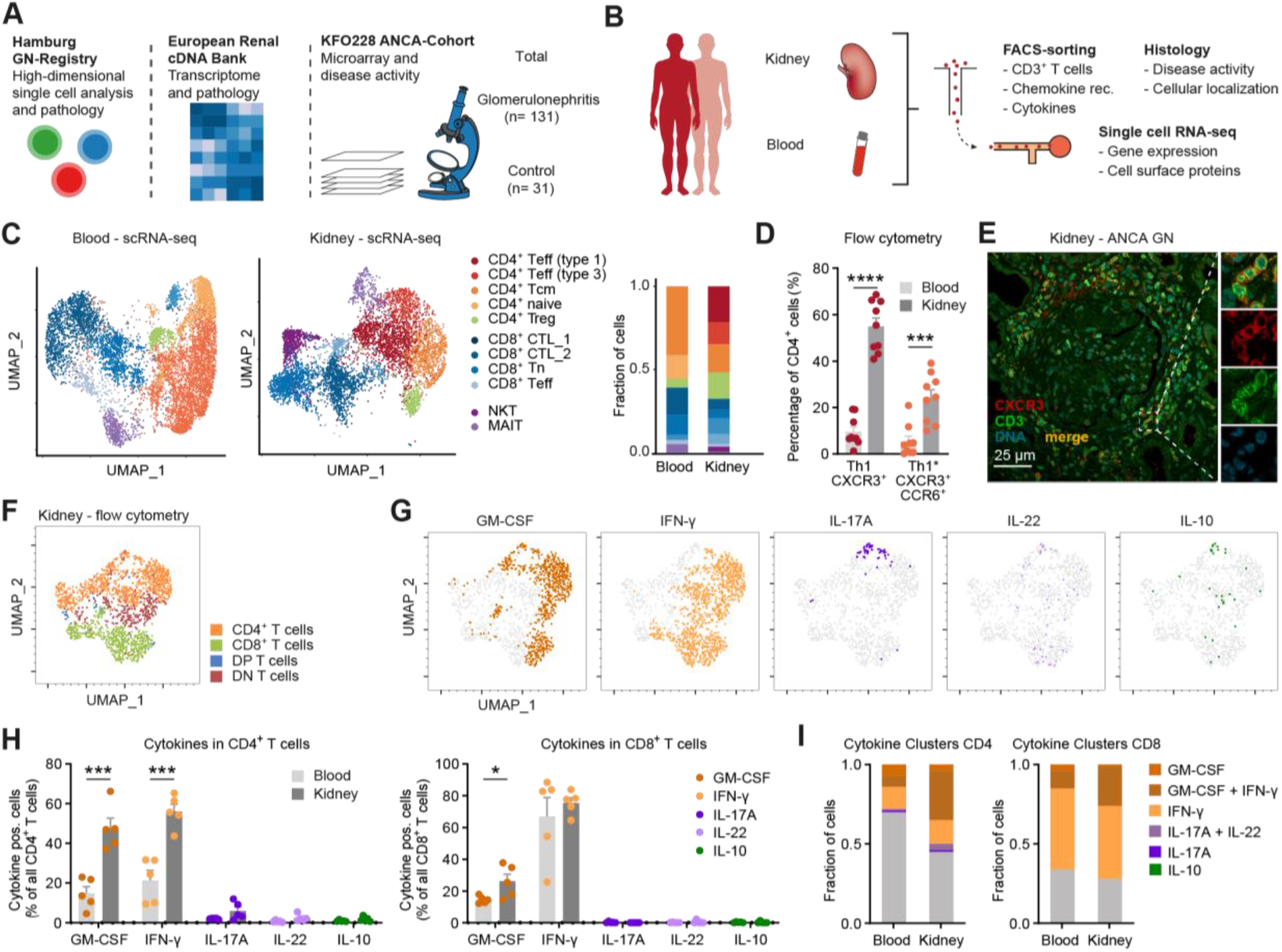
Identification of GM-CSF-producing T cells in patients with crescentic glomerulonephritis. **(A)** Overview of three cohorts used to assess the cellular immune response in patients with crescentic glomerulonephritis. **(B)** Handling of kidney tissue specimens and peripheral blood samples for high-dimensional profiling by FACS (n=14), scRNA-seq (n=8), and immunofluorescence (n=20). **(C)** scRNA seq of FACS-sorted CD3^+^ T cells from peripheral blood and the kidney of patients with ANCA-GN were processed for unsupervised clustering and UMAP, as well as relative contribution of different clusters. **(D)** FACS-based quantification of chemokine receptor expression from peripheral blood and kidney T cells of patients with ANCA-GN (n=9). **(E)** Representative immunofluorescence staining of renal CXCR3^+^ T cells in a patient. **(F-I)** Peripheral blood and kidney T cells of patients with ANCA-GN were stimulated with PMA/ionomycin and cytokine production was analyzed by FACS. **(F)** UMAP of FACS analyzed T cells and manually gated population identification as indicated (n=5). **(G)** Cytokine positive cells in the same UMAP. **(H)** Frequencies of cytokine^+^ cells in peripheral blood and kidney CD4^+^ and CD8^+^ T cells. **(I)** Cluster frequencies of type 1 cytokine (GM-CSF and IFN-γ), type 3 cytokine (IL-17 and IL-22) and IL-10 expressing CD4^+^ and CD8^+^ T cells (* P<0.05, *** P<0.005, **** P<0.001).

### GM-CSF-producing Th1 cells accumulate in the kidney of ANCA-GN patients

To capture the peripheral blood and kidney T cell landscape of crescentic glomerulonephritis, we performed scRNA sequencing of CD3^+^ T cells sorted from pair matched blood and renal biopsy samples of patients with ANCA-associated glomerulonephritis (ANCA-GN) (Figure 1B). Unsupervised clustering recognized 8 clusters in the blood and 11 clusters in the inflamed kidney. Based on the transcriptional profile of individual cells, we identified the following T cell subsets: naïve, central memory, type 1 / type 3 effector and regulatory CD4^+^ T cells; naïve, cytotoxic and tissue-resident memory CD8^+^ T cells; mucosal associated invariant T cells (MAIT) and natural killer T cells (NKT) (Figure 1C, Supplemental Figure 1A and B). In particular, type 1 (Th1) and type 3 (Th17) effector T cells were found enriched in the kidney of patients with ANCA-GN. The dominance of Th1 cells in the kidney was also confirmed by flow cytometry for the chemokine receptors CXCR3 and CCR6 to identify Th1 and Th1* cells (Figure 1D and Supplemental Figure 2A) (*15*). Of note, CXCR3 immunofluorescence staining revealed that kidney-invading Th1 cells precisely localized to the inflamed glomerulus, the periglomerular infiltrate and the tubulointerstitial compartment in ANCA-GN patients (Figure 1E). Flow cytometry-based cytokine staining of polyclonal stimulated blood and kidney leukocytes from ANCA-GN patients highlighted the prevalence of GM-CSF and IFN-γ producing cells among CD4^+^ and CD8^+^ T cells, while IL-17A producing cells were less frequent, and IL-22 and IL-10 were hardly detectable (Figure 1F, 1G, and Supplemental Figure 1C and D), as well as their selective enrichment in the inflamed kidney compared to the blood (Figure 1H and 1I). Altogether, our results show that kidneys from ANCA-GN patients show prominent accumulation of GM-CSF producing Th1 cells that localize at the main sites of tissue destruction.

### CD4^+^ T cell-derived GM-CSF drives experimental GN

To investigate the functional role of GM-CSF in kidney inflammation, we used a well-characterized preclinical mouse model of crescentic GN (cGN) that recapitulates the human disease in terms of functional impairment, tissue injury and immune cell infiltration (*4, 16, 17*). In a first step, we analyzed cGN-associated tissue damage in wild-type and GM-CSF-deficient mice (Figure 2A). The evaluation of kidney sections revealed a significant reduction of glomerular crescents and tubulointerstitial injury in GM-CSF-deficient mice (Figure 2B and 2C). Moreover, blood urea nitrogen (BUN) levels, which are a measure of functional kidney impairment, were significantly reduced in GM-CSF-deficient compared to wild-type cGN mice (Figure 2C). Immunohistochemistry showed that renal infiltration of F4/80^+^ mononuclear phagocytes was reduced in *Csf*2^−/−^ mice, while recruitment of CD3^+^ T cells remained unchanged (Supplemental Figure 3A).

**Figure 2:**
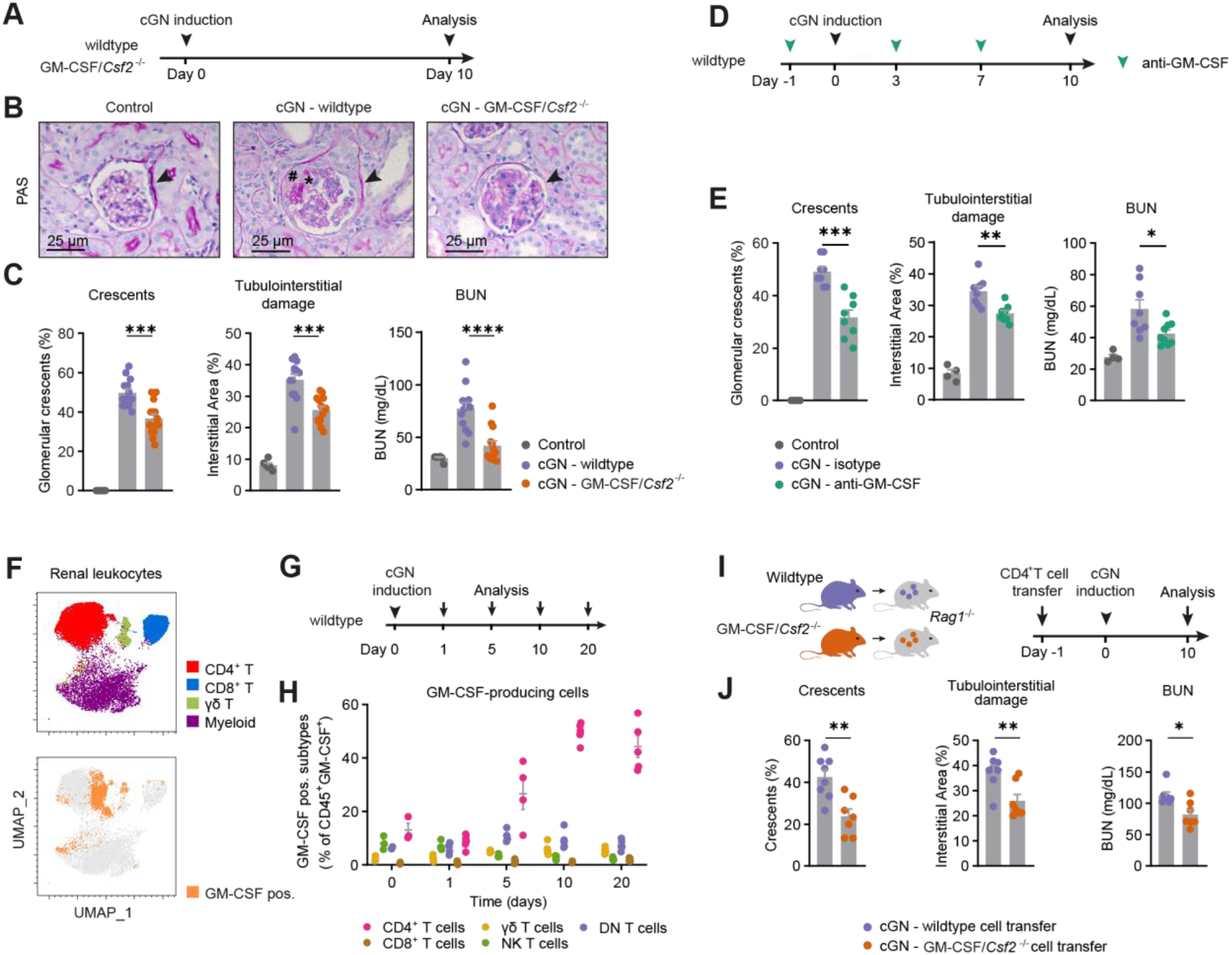
T cell-derived GM-CSF drives renal tissue damage in crescentic GN (cGN). **(A-C)** cGN induction in wildtype and GM-CSF/*Csf2*^-/-^ mice. **(B)** Representative photographs of PAS-stained kidney sections from control, nephritic wild-type, and nephritic GM-CSF-deficient mice 10 days after cGN induction. **(C)** Quantification of glomerular crescent formation, tubulointerstitial damage, and BUN levels. (**D-E**) Neutralization of GM-CSF using a monoclonal antibody (mAb). **(D)** cGN was induced in wildtype mice and anti-GM-CSF was injected at indicated time points. **(E)** Quantification of glomerular crescents, tubulointerstitial damage and BUN levels. **(F-H)** Cellular source of GM-CSF. **(F)** Dimensional reduction of FACS data by UMAP and manual gating of renal leukocytes from nephritic wild-type mice ten days after cGN induction. **(G)** cGN was induced in wildtype mice and analyzed at indicated time points. **(H)** Quantification of leukocyte subset contribution to renal GM-CSF production in the course of cGN. **(I-J)** Adoptive CD4^+^ T cell-transfer into *Rag1*^−/−^ mice and induction of cGN. **(J)** Quantification of glomerular crescents, tubulointerstitial damage, and BUN levels. Symbols represent individual data points with the mean as a bar (* P<0.05, ** P<0.01, *** P<0.005, **** P<0.001). Data are representative of two independent experiments (► Bowman’s capsule, * glomerular necrosis, # glomerular crescent).

To analyze the effect of interventional GM-CSF targeting, nephritic wild-type mice were treated with either an anti-GM-CSF-neutralizing antibody or an isotype control antibody (Figure 2D). Anti-GM-CSF treatment ameliorated cGN in terms of renal tissue injury and BUN levels (Figure 2E and Supplemental Figure 3B).

To identify the cellular source of GM-CSF, we performed time course analyses in nephritic mice (days 0 to 20). In the course of cGN, renal GM-CSF mRNA and protein expression peaked around day 10, the time point with the largest CD4^+^ T cell infiltrate (*18*), whereas in the plasma, GM-CSF was detectable at lower levels and peaked at day 5 (Supplemental Figure 3D-F). Flow cytometric analysis after stimulation with PMA/ionomycin revealed that CD4^+^ T cells were the major cellular source of GM-CSF at day 10, but its production was also observed to a lesser degree in CD8^+^, CD4^−^CD8^−^ (double negative) T cells, CD3^+^ NK1.1^+^ natural killer T cells, and γδ T cells throughout the disease course (Figure 2F-H).

To determine whether CD4^+^ T cell-derived GM-CSF is sufficient to drive renal tissue injury, we performed adoptive CD4^+^ T cell transfer of from either wild-type or GM-CSF-deficient donor mice into *Rag1*^−/−^ mice, which lack T and B cells, and induced cGN (Figure 2I). Indeed, selective GM-CSF-deficiency in CD4^+^ T cells resulted in an ameliorated course of cGN (Figure 2J and Supplemental Figure 3G). In conclusion, our data show that CD4^+^ T cell-derived GM-CSF is a major driver of renal tissue injury in cGN and that antibody-based GM-CSF neutralization is effective in restraining cGN immunopathology.

### Monocyte-derived Cells (MdCs) are the major target cells of GM-CSF

Next, we sought to dissect the effects of GM-CSF signaling on the immune cell composition in the kidney. To this end, we compared CD45^+^ cells from wild-type and GM-CSF-deficient nephritic mice by high-dimensional flow cytometry (spectral FACS) and scRNA-seq in two independent sets of experiments (Figure 3A and 3B). Both analyses pointed to a population of monocyte-derived cells (MdCs; CD45^+^CD11b^+^CD11c^+^CD88^+^CD64^+^F4/80^+^MHCII^+^ cells) that was more abundant in the inflamed kidney of wild-type compared to GM-CSF/*Csf*2^−/−^ mice, indicating that this population could be dependent on GM-CSF signaling (Figure 3A, 3B and Supplemental Figure 4A-C). Of note, renal MdC numbers were low under homeostatic conditions (Supplemental Figure 4A), supporting their origin from circulating monocytes that infiltrate the nephritic kidney. In line with this, MdCs represent a highly inflammatory mononuclear phagocyte (*19–21*) and MdCs from the inflamed kidney (our dataset) displayed high expression of the GM-CSF receptor (*CSF2Ra/b*; Supplemental Figure 4D). Taken together, this prompted us to hypothesize that MdCs could mediate GM-CSF-induced renal tissue damage in cGN.

**Figure 3:**
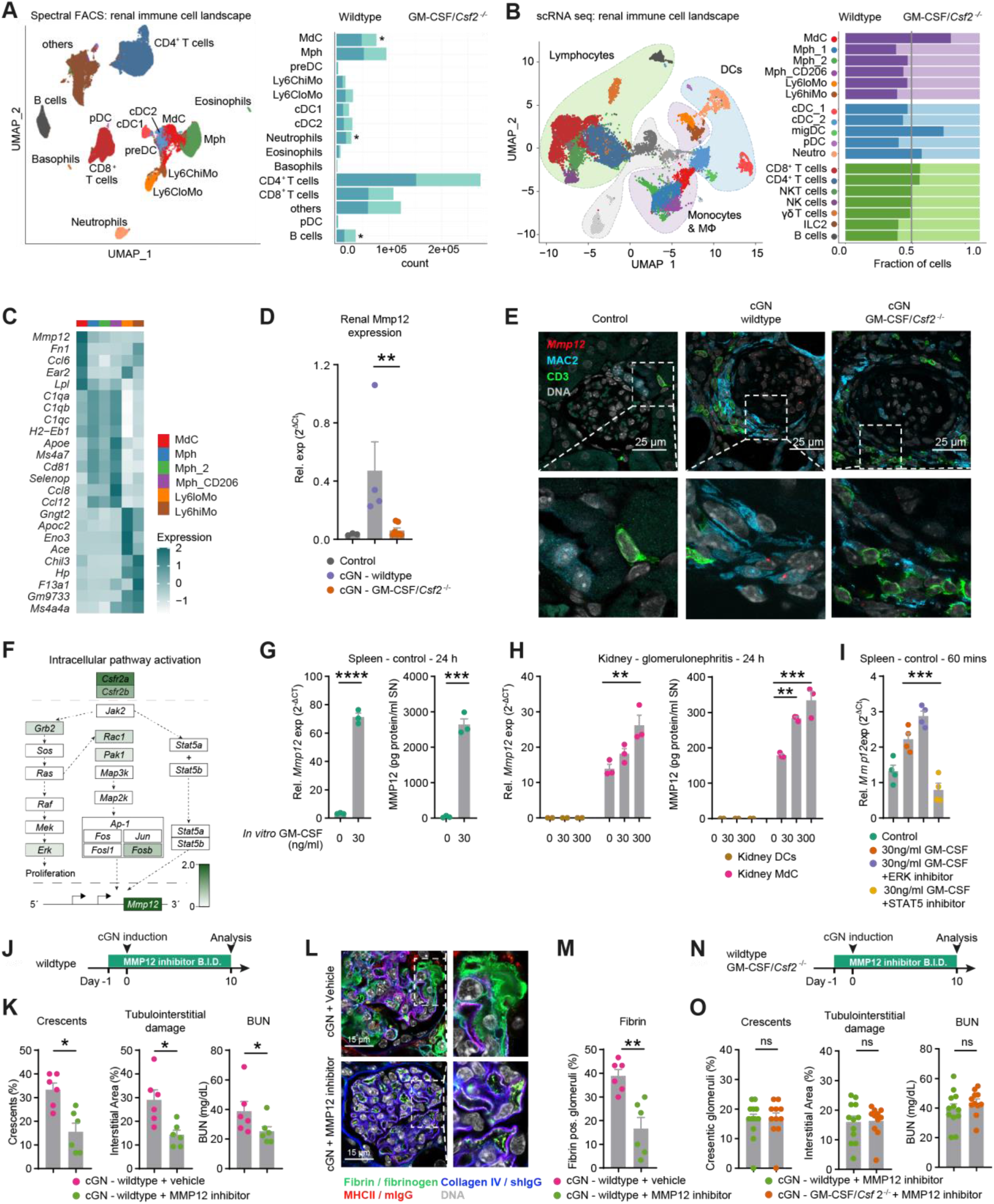
GM-CSF induces renal immunopathology via MMP12 expression in monocyte derived cells. **(A)** Unsupervised clustering and UMAP of FACS-analysis of renal leukocytes from nephritic wild-type and GM-CSF/*Cf2*^−/−^ mice. Cell counts in the respective clusters from wildtype and GM-CSF/*Csf2*^-/-^ animals. **(B)** Unsupervised clustering and UMAP dimensional reduction of scRNA-seq of renal leukocytes from wildtype and GM-CSF/*Csf2*^−/−^ mice ten days after cGN induction. Clusters were annotated according to their gene expression profiles. Fraction of cells in the respective clusters from wildtype and GM-CSF/*Cf2*^−/−^ animals. **(C)** Heatmap of key marker gene expression of the indicated clusters form scRNA-seq. **(D)** RT-PCR analysis of renal *Mmp12* expression in wildtype and GM-CSF^−/−^ mice. **(E)** Combined immunofluorescence staining of CD3^+^ (green), MAC2^+^ (turquoise), and FISH (*Mmp12*, red) of renal leukocytes in wildtype and GM-CSF/*Csf2*^−/−^ mice. **(F)** Pathway analysis of differentially expressed genes of renal MdCs from nephritic wildtype and GM-CSF/*Csf2*^−/−^ mice (color code indicating the log2-fold change in wildtype versus GM-CSF/*Csf2*^−/−^ animals). **(G)** FACS sorted MdC like cells from the spleen of control mice were *in vitro* cultured for 24 hours with/without addition of recombinant mouse GM-CSF. *Mmp12* gene expression and protein expression in the supernatant were quantified. **(H)** FACS sorted MdCs from kidneys of cGN mice were *in vitro* cultured for 24 hours with/without addition of recombinant mouse GM-CSF. *Mmp12* gene expression and protein expression in the supernatant were quantified. **(I)** FACS sorted MdC like cells from spleens of control mice were *in vitro* cultured for 1 hour with/without addition of recombinant mouse GM-CSF and with addition of indicated pharmacological inhibitors. *Mmp12* gene expression was quantified by RT-PCR. **(J-M)** Crescentic GN induction and treatment with pharmacological MMP12 inhibitor or vehicle. **(J)** cGN was induced in wildtype mice and MMP12 inhibitor was given by oral gavage at indicated time points (bis in die, B.I.D.). **(K)** Quantification of glomerular crescents, tubulointerstitial damage, and BUN levels. **(L)** Representative confocal micrographs depicting extent of glomerular injury in kidneys of mice with/without MMP12 inhibitor treatment. Note the strong fibrin/fibrinogen signal within the capillaries of the cGN + vehicle glomerulus, whereas msIgG and shIgG/CollagenIV signals at the glomerular filtration barrier are comparable between both groups. **(M)** Quantification of fibrin positive glomeruli detected by immunohistochemistry. **(N)** cGN was induced in wildtype mice and GM-CSF-deficient mice as well as MMP12 inhibitor was given by oral gavage at indicated time points (bis in die, B.I.D.). **(O)** Quantification of glomerular crescents, tubulointerstitial damage, and BUN levels. Symbols represent individual data points with the mean as a bar (* P<0.05, ** P<0.01, *** P<0.005, **** p<0.001). Data are representative of two independent experiments.

### GM-CSF drives MMP12 expression in MdCs

Next, we aimed to identify molecular pathways that might mediate the MdC effector function in kidney immunopathology. An analysis of cluster-defining genes from the kidney scRNAseq data set from mice with cGN identified matrix metalloproteinase-12 (*Mmp-12*) as the most differentially expressed gene in MdCs as compared to the other myeloid clusters (Figure 3C), and *Mmp12* expression was dependent on GM-CSF in the inflamed kidney (Figure 3D). Matrix metalloproteinases are involved in the breakdown of extracellular matrices in physiological processes and disease. MMP12 particularly targets elastin and type IV collagen (*22*), which are key components of the interstitial matrix and the basement membrane in the kidney. MMP12 could, indeed, contribute to the processes involved in glomerular crescent formation and necrosis (*23*). Of note, immunofluorescence staining in combination with fluorescence *in-situ* hybridization (FISH) revealed the adjacent localization of CD3^+^ T cells and *Mmp12*-expressing MAC2^+^ myeloid cells to the periglomerular area (Figure 3E). *Mmp12* expression was only detected in MAC2^+^ myeloid cells of wild-type cGN, but not in the kidney of nephritic GM-CSF-deficient mice (Figure 3E).

To determine the GM-CSF downstream signaling in MdCs, we performed pathway analysis based on GO terms using differentially expressed genes in MdCs from wild-type versus GM-CSF-deficient nephritic mice. These analyses revealed the activation of Jak2 downstream signaling. Differentially expressed genes in our dataset are indicated in the intracellular pathway scheme (Figure 3F). Concordantly, GM-CSF directly induces MMP12 expression in sorted MdC-like spleen and kidney cells, but not in DCs (Figure 3G-H). Moreover, inhibition of the JAK2/STAT5 pathway abolished GM-CSF induced MMP12 expression, in contrast to ERK inhibition (Figure 3I). Together, GM-CSF control MMP12 expression in MdCs *in vivo* and *in vitro* via JAK2/STAT5 signaling.

### The GM-CSF – MdC-derived MMP-12 axis drives kidney injury

In view of the GM-CSF pathogenic role in human and experimental GN and the high inflammatory potential of MMP12-expressing MdCs, we next investigated as to whether MMP12 represents an effector to immune-mediated kidney damage. Therefore, we blocked MMP12 enzymatic activity via treatment with the specific inhibitor MMP408 during cGN course (days −1 to 10) (*24*) (Figure 3J). Upon induction of cGN, MMP12 inhibition reduced renal pathology compared to vehicle-treatment (Figure 3K and Supplemental Figure 5A). Moreover, we observed significant reduction of glomerular necrosis and destruction of glomerular capillary wall in MMP12 inhibitor-treated mice (Figure 3L and M).

Importantly, MMP12 inhibition abolished the pathological and clinical differences between nephritic wild-type and GM-CSF-deficient mice, indicating that the ameliorated cGN in GM-*CSF*/*CSF2*^−/−^ mice is a consequence of the neutralized GM-CSF – MMP12 pathway (Figure 3N, 3O and Supplemental Figure 5B). Taken together, GM-CSF-dependent MMP12 activity in MdCs drives immunopathology in cGN.

### GM-CSF-transcriptional signature is upregulated in patients with GN and correlate with the degree of renal pathology

We next addressed the question whether GM-CSF-producing T cells in the kidneys of GN, and subsequent *MMP12* expression, might have translational value for patients with GN. To this end, we interrogated the bulk RNA dataset of the European Renal cDNA Bank database, including renal biopsies of patients with ANCA-GN, lupus nephritis (caused by systemic lupus erythematosus), and IgA nephropathy (IgAN) as well as control biopsies from living donor kidney transplantation, for the expression levels of 25 GM-CSF-signature genes (*GM-CSF tissue score*) (13). *GM-CSF tissue score* was up-regulated in all GN groups compared to controls, but had the highest value in the kidney of patients with ANCA-GN (Figure 4A and 4B). This might reflect the more severe form of crescentic GN in this disease group when compared to lupus nephritis and IgA nephropathy. Of note, the *GM-CSF tissue score* was found to directly correlate to the severity of renal tissue damage in terms of glomerular necrosis and crescent formation in ANCA-GN patients (Figure 4C and 4D).

**Figure 4:**
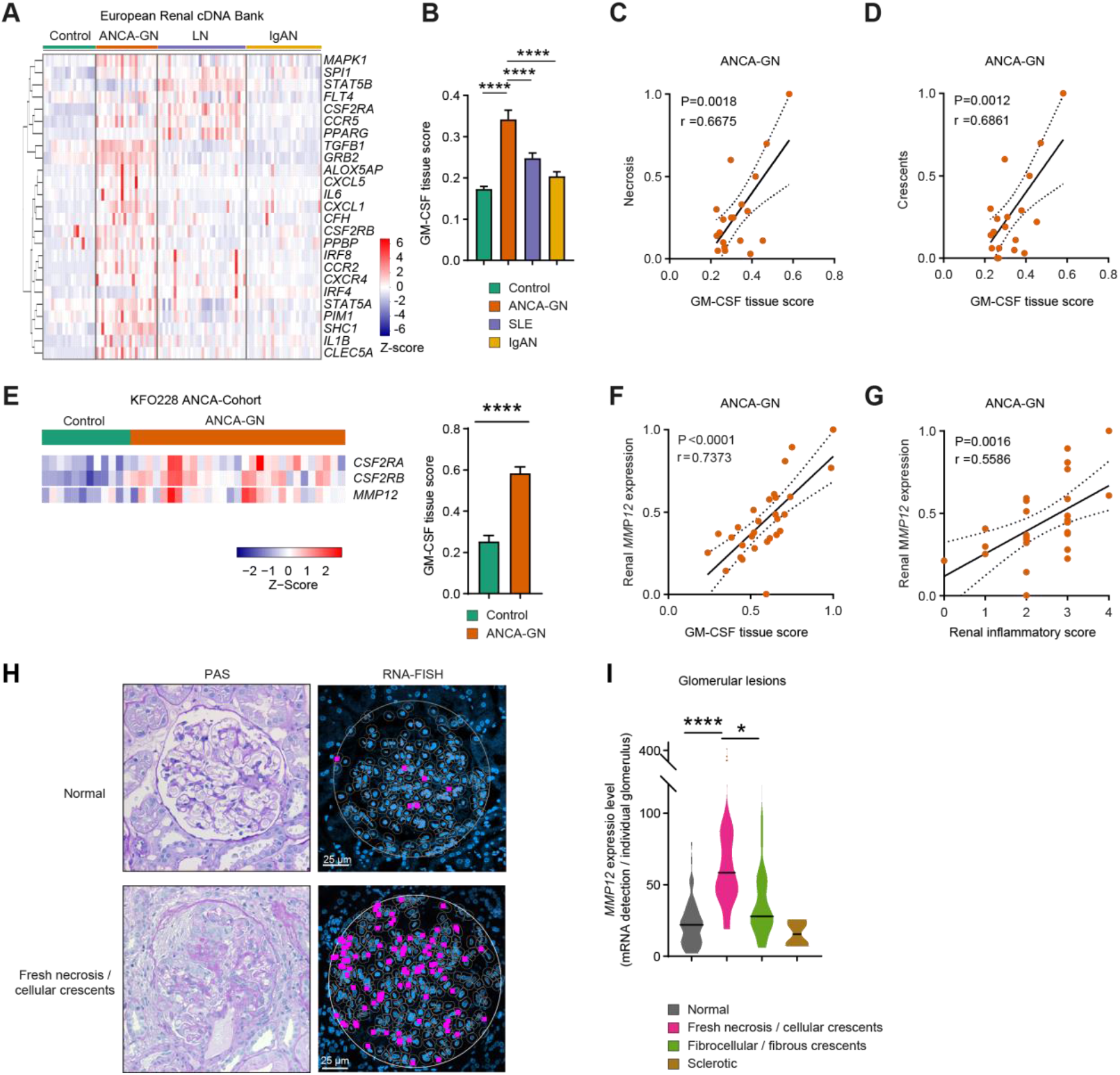
Enrichment of GM-CSF signaling and MMP12-expressing myeloid cells in renal biopsies from patients with glomerulonephritis. **(A)** Gene expression of GM-CSF signaling related genes (tissue score) in glomerular kidney samples of controls, ANCA-GN, SLE/Lupus Nephritis and IgA nephropathy patients from the European Renal cDNA Biobank (ERCB). **(B)** Quantification of the GM-CSF tissue score based on cumulative gene expression in the respective patient groups. **(C-D)** Correlation of **(C)** necrosis and **(D)** percentage of glomeruli with crescents and GM-CSF tissue score. **(E)** Renal GM-CSF receptor (*CSF2RA* and *CSF2RB*) and *MMP12* tissue gene expression in controls and ANCA-GN patients from the CRU 228 ANCA-Cohort. **(F)** Correlation of renal MMP12 expression with the GM-CSF tissue score or **(G)** renal inflammation score in renal biopsies of patients with ANCA-GN. **(H)** Representative image of *MMP12* FISH staining and corresponding PAS staining of the human kidney biopsy, blue (cell nucleus), grey (cellular dimension), magenta (*MMP12*). **(I)** Quantification of *MMP12* signals by image-based software (ACD QuPath) in normal glomeruli and glomeruli with fresh necrosis and/or cellular crescents, with fibrocellular and fibrous crescents as well as sclerotic glomeruli. Symbols represent individual data points with the mean as a bar (* P<0.05, ** P<0.01, *** P<0.005, **** P<0.001).

Validation analysis in an independent ANCA-GN cohort (CRU 228 ANCA Cohort) (*14*) not only confirmed the upregulation of GM-CSF–associated genes in the kidney of the ANCA-GN patients (Figure 4D), but also showed its direct correlation with renal *MMP12* expression (Figure 4F). Moreover, the degree of renal *MMP12* expression correlated with the renal inflammatory activity (Figure 4G). Consistently, combined FISH and immunofluorescence analyses in renal biopsy specimens from patients with ANCA-associated GN demonstrated that *MMP12*-expressing myeloid cells co-localized with CD3^+^ T cells to areas of highly active and destructive tissue inflammation (Supplemental Figure 6A and 6B). FISH analysis of 160 individual glomeruli from 20 ANCA patients of the Hamburg GN Registry revealed that *MMP12* expression was highest in glomeruli with fresh necrosis and/or cellular crescents, significantly lower in glomeruli with fibrous cellular and fibrous crescents, and hardly detectable in sclerotic and normal glomeruli (Figure 4H and 4I).

## DISCUSSION

Kidney-invading T cells promote tissue injury and loss of renal function in human and experimental crescentic glomerulonephritis (*5, 8, 25*). Here, we identified the underpinning mechanism of how T cells, through the release of GM-CSF, license highly proinflammatory monocyte-derived cells to cause renal immunopathology in both pre-clinical mouse models and in patients with crescentic glomerulonephritis.

GM-CSF was originally identified as a hematopoietic growth factor because of its ability to promote the survival and activation of macrophages and granulocytes as well as dendritic cells from precursor cells (*26*). *In vivo*, in the steady state GM-CSF is required only for development and maintenance of alveolar macrophages in the mammalian lungs (*27*). In inflammation however, GM-CSF prediction is highly elevated and plays a central role in connecting the adaptive and innate parts of the immune system by acting directly on mature myeloid cells. Myeloid cells sense GM-CSF *via* specific GM-CSF receptors, resulting in a pro-inflammatory phenotype with increased effector functions (*28, 29*). In the context of chronic immune-mediated or infectious diseases, GM-CSF might have significant deleterious effects by promoting tissue hyperinflammation and injury (*30*). The pathological effects of GM-CSF are best characterized in experimental autoimmune encephalomyelitis (EAE), a model of multiple sclerosis (MS). Here, GM-CSF is primarily produced by CD4^+^ T cells (*31*) and has a key function in driving destructive neuroinflammation (*20, 31–34*). In the field of immune-mediated kidney disease it was shown almost two decades ago that GM-CSF deficient mice develop less severe glomerulonephritis but the underlying mechanism remained unclear (*35, 36*).

Our characterization of the renal T cell infiltrate in human and experimental cGN, identified GM-CSF^+^ T cells as a major driver of kidney immunopathology and Th1 subsets as the major source of this cytokine in the inflamed tissue. The differential analysis at single cell level of cGN immune landscape in wildtype versus *Csf*2^−/−^ mice indicated MdCs as the major target cells of GM-CSF in renal inflammation. MdCs have been proposed to differentiate in the inflamed tissue from infiltrating Ly6C^hi^ monocytes (*19, 37*). The heterogenous MdC population with features of both macrophages and DCs might act as a highly proinflammatory phagocytes promoting tissue inflammation and damage via multiple effector mechanism encompassing ROS, RNS and inflammatory cytokine production (*20, 21, 32*). Our present study uncovered a previously unknown GM-CSF-induced effector mechanism of MdCs.

Of note, kidney MdCs are characterized by high expression of MMP12, a member of the matrix metalloproteinase family of zinc-dependent proteolytic enzymes, which are inflammatory mediators that might degrade key components of the extracellular matrix in the kidney, such as collagen type IV and elastin (*22, 38*). We found that T cell-derived GM-CSF directly activates *MMP12* expression by MdCs via the GM-CSF receptor – JAK/STAT5 signaling pathway. Pharmacological MMP12 neutralization resulted in less severe glomerulonephritis in terms of glomerular crescent formation, tubulointerstitial injury, and renal function. In line with this, GM-CSF tissue signaling and *MMP12* expression correlated with the severity of renal inflammation in patients with ANCA-GN, and *MMP12* expressing myeloid cells were localized to areas of severe and destructive glomerular injury. This indicates that MMP12 degrades key structural components of the kidney such as the glomerular basement membrane and the Bowman’s capsule and thereby significantly contributes to kidney pathology.

We could also identify a distinct cluster of MMP12-expressing myeloid cells in the lungs of patients with severe COVID-19 (Supplemental Figure 7A-C) together with up-regulated GM-CSF expression (*39*). This is of particular interest because the basement membrane in the kidney and the lung have a similar composition (*40*) and both can be targeted by GM-CSF-induced MMP12. This suggests a more general function of this pathway across different inflammatory diseases and tissues. Overall, our study provides a new mechanistic basis for the immunopathology of T cell-mediated diseases, identifies the GM-CSF / MMP12 network as a promising therapeutic target in T cell–driven inflammatory diseases and provides a rationale for an anti-GM-CSF clinical trial in patients with crescentic glomerulonephritis.

## METHODS

### Human studies

Human kidney sections and clinical parameters were analyzed from patients with glomerulonephritis included in the Hamburg GN Registry (cohort 1), the European Renal cDNA Bank (cohort 2) and the Clinical Research Unit 228 (CRU 228) ANCA cohort (cohort 3). For some of the analyses, matched blood samples from the same patients were used. For the analysis of cells from control samples, kidneys undergoing living donor transplantation were used. Cells from patients with severe COVID-19 were obtained from bronchioalveolar lavage fluid (BALF) as previously published (*39*). These studies were approved by the *Ethik-Kommission der Ärztekammer Hamburg*, (local ethics committee of the chamber of physicians in Hamburg) and were conducted in accordance with the ethical principles stated in the Declaration of Helsinki. Informed consent was obtained from all participating patients. Information on the patient cohorts are provided in the Supplemental Table 1.

### Animal experiments

Mouse experiments were carried out in accordance with the national guidelines. The protocols were approved by local ethics committees. Mice were housed under specific pathogen free conditions. All experiments were conducted with age matched male mice on a C57BL/6 background. The following genetically modified mouse strains were used: *Csf2*^−/−^ mice B6.129S-Csf2tm1Mlg/J and *Rag1*^−/−^ mice B6.129S7-Rag1tm1Mom/J (both Jackson Laboratory). Experimental crescentic glomerulonephritis was induced in male mice by i.p. application of sheep serum directed against the glomerular basement membrane (2.5 mg/g bodyweight), as previously described (*12*).

GM-CSF was neutralized by i.p. injection of 300 μg of anti-GM-CSF (clone MP1-22E9, BioXCell, Lebanon, NH) in 200 μl InVivoPure pH 7.0 Dilution Buffer (BioXCell) on days −1, 3, and 7 after induction of crescentic GN. Control mice received InVivoMAb rat IgG2a isotype control (clone 2A3, BioXCell). Pharmacological inhibitor of MMP12 (MMP408, Calbiochem, San Diego, CA) was administered by oral gavage of 0.25 ml per mouse at 5 mg/kg and twice per day. The compound was put into solution using compound vehicles, 2% Tween 80 and 0.5% methylcellulose.

### Reconstitution of *Rag1*^−/−^ mice with CD4^+^ T cells

For adoptive cell transfer, lymphocytes were isolated from spleens of naïve C57BL/6 or GM-*CSF*/*CSF2*^−/−^ mice and CD4^+^ T cells were enriched by magnetic cell sorting (Mouse CD4^+^ T Cell Isolation Kit, Miltenyi Biotec, Bergisch Gladbach, Germany). Enrichment of CD4^+^ T cells was verified by flow cytometry and consistently reached >97%. *Rag1*^−/−^ mice were reconstituted i.v. with 5×10^5^ CD4^+^ T cells per animal (*41*).

### Histopathology, immunohistochemistry and immunofluorescence

Glomerular crescent formation was assessed in a blinded fashion in 30 glomeruli per mouse in periodic acid-Schiff (PAS)-stained paraffin sections (*11*), and tubulointerstitial injury was assessed as described (*17,18*). For immunohistochemistry, staining of paraffin-embedded kidney sections (2 μm) using the following antibodies was performed as previously described (*18*): CD3 (A0452, Dako, Germany), F4/80 (BM8; BMA, Hamburg, Germany). For immunofluorescence, the following primary antibodies were incubated overnight in blocking buffer at 4°C: CD68 (KP1, Invitrogen, Invitrogen, Carsbad, CA), CD3 (A0452, Dako, Glostrup, Denmark), CXCR3 (1C6, BD Biosciences), Mac-2 (M3/38, Cedarlane, Burlington, Canada), CXCR3 (Bioss Antibodies, Woburn, MA), MHCII (sc-59322, Santa Cruz), Fibrin/Fibrinogen (A0080, Dako), CollagenIV (1340-01, SouthernBiotech). After washing in PBS, fluorochrome-labeled secondary antibodies and Hoechst (Molecular Probes, Eugine, OR) were applied. Staining was visualized using an LSM800 with Airyscan and the ZenBlue software, or with an ELYRA PS.1 SIM microscope and the ZenBlack software (all Zeiss). *Mmp12* mRNA detection (FISH) in mice and human kidney sections was carried out using RNAscope Multiplex Fluorescent Assay (412658 (human) or 406558 (mouse), Advanced Cell Diagnostics, Newark, CA). The slides were imaged using a Zeiss LSM800 confocal microscope and analyzed with ZEN software (Carl Zeiss, Jena, Germany). For quantification of MMP12 expression the ACD QuPath software was used.

Renal inflammatory activity score: Combined histopathological score of the following four parameters: Percentage of cellular crescents / glomerular necrosis (0 <50%, 1 >50%); percentage of glomerular sclerosis (0 >50%, 1 <50%); percentage of interstitial inflammation / cell infiltration (0 <25%, 1 >25%); presence of a periglomerular cell infiltrate or a Bowman’s capsule rupture (0 <25%, 1 >25%). The maximum renal inflammatory activity score is four. Individual glomeruli of ANCA-GN patients were categorized as normal, as glomeruli with fresh necrosis and/or cellular crescents, as glomeruli with fibrocellular and fibrous crescents or as sclerotic glomeruli, according to the Renal Pathology Society (*42*).

### Isolation and flow cytometric analysis of human and murine leukocytes

Single cell suspensions were obtained from human kidney biopsies by enzymatic digestion in RPMI 1640 medium with collagenase D at 0.4 mg/ml (Roche, Mannheim, Germany) and DNase I (10 μg/ml, Sigma-Aldrich, Saint Louis, MO) at 37° C for 30 minutes followed by dissociation with gentleMACS (Miltenyi Biotec). Leukocytes from blood samples were separated using Leucosep tubes (Greiner Bio-One, Kremsmünster, Austria). Samples were filtered through a 30 μm filter (Partec, Görlitz, Germany) before antibody staining and flow cytometry.

Cells from murine spleens were isolated by squashing the organ through a 70 μm cell strainer. Erythrocytes were lysed using a lysis buffer (155 mM NH4Cl, 10 mM KHCO3, 10 μM EDTA, pH 7.2). Kidneys were enzymatically digested with 400 μg/ml collagenase D (Roche) and 10 U/ml DNase I (Sigma-Aldrich) for 45 min at 37°C. Subsequently, leukocytes were isolated by density gradient centrifugation using 37% Easycoll (Merck Millipore) and a filtration step using a 30 μm cell strainer (Partec).

Previously described methods for immune-cells isolation and for cytokines detection from murine kidneys and spleens were used (*12*). Cells were surface stained with fluorochrome-conjugated antibodies (human: CD45 (H130), CD3 (OKT3), γδ-TCR (B1), CD4 (RPA-T4), CD8 (RPA-T8), CD14 (HCD14), CD16 (3G8), CD19 (HIB19), CD25 (BC96), CD69 (FN50), CD103 (Ber-ACT8 (BD), CD45RA (HI100), CD11b (M1/70), CD11c (42981), HLA-DR (G46-6), GM-CSF (VD2-21C11), IL-10 (JES3-19F), IL-17A (BL168), IL-17F (SHLR17), IL-22 (G12A41), IFN-γ (4S.B3), TNF-α (MAB11), CXCR3/CD183 (Go25H7), CCR4/CD194 (L291H4), CCR6/CD196 (Go34E3), CCR7/CD197 (Go43H7), CCR10/CD (6588-5); mouse: CD45 (30-F11), CD3 (145-2C11), γδ-TCR (eBio GL3), CD4 (RM4-5), CD8 (53-6.7), CD19 (6D5), CD25 (PC61), CD11b (M1/70), CD11c (N418/HL3), CD64 (X54-5/7.1), Ly6C (AL-21/HK1.4), Ly6G (1A8), F4/80 (BM8), MHCII (M5/114.15.2), NK1.1 (PK136), CD206 (CO68C2), GM-CSF (MP1-22E9), IL-10 (JES5-16E3), IL-17A (bd TC11-18H10), IL-17F (9D3.1C8), IL-22 (poly 5164), IFN-γ (XMG1.2), TNF-α (MP6-XT22), Arg-1 (A1exF5), CX3CR1 (SA011F11), Tim4 (RMT4-54), CD206 (CO68C2), CD44 (IM7), XCR1 (ZET), CD16/32 (93), CD88 (20/70), NOS2 (W16030C), PDL1 (10F.9G2), CD16.2 (9E9), CD38 (90); all BD Biosciences, Biolegend or eBiosciences) and a fixable dead cell stain (Molecular Probes) to exclude dead cells from analysis. For intracellular staining, samples were processed using Cytofix/Cytoperm (BD Biosciences) according to the manufacturer’s instructions.

### Flow cytometry and cell sorting

Samples were measured with a LSR II and Symphony A3 (both BD Biosciences) or Cytek Aurora (Cytek Biosciences, Amsterdam, The Netherlands). Data analysis was performed using the FlowJo software (Treestar, Ashland, OR) or the FACSDiva software (BD Biosciences). FACS-sorting was performed on a AriaFusion or AriaIIIu (BD Biosciences).

### *In vitro* stimulation of cells

MdC like cells (CD45^+^CD3^-^CD11b^+^CD64^+^) from the spleen and kidney of mice were purified using anti-FITC MicroBeads (Miltenyi Biotec) and FACS sorted. 1 ×10^4^ MdCs were *in vitro* cultured in a volume of 200 μl of RPMI 1640 medium containing 10% FCS, streptomycin and penicillin. Cells were treated with 30 ng/ml recombinant mouse GM-CSF (Biolegend) with/without selective ERK inhibitor (Tocris, Bristol, UK) or selective STAT5 inhibitor (Sigma-Aldrich). After 20 minutes (short term) or 1 day (long term), the expression of *Mmp12* was determined by RT-PCR or in the supernatants by ELISA (Abcam, Cambridge, UK).

### Single cell RNA sequencing

To perform scRNA-seq, single cell suspensions were obtained from human and mouse kidney samples. Cell hashing was performed according to manufacturer’s instructions (Biolegend). FACS-sorted CD45^+^ cells or CD3^+^ T cells were used for droplet-based single cell analysis and transcriptome library preparation using the Chromium Single Cell 5′ Reagent Kits v2 according to the manufacture’s protocols (10× Genomics, Pleasanton, CA). The generated scRNA-seq libraries were sequenced on a NovaSeq6000 system (100 cycles) (Illumina, San Diego, CA).

### Alignment, quality control and pre-processing of single-cell RNA-sequencing data

Quality control and scRNA-seq pre-processing was performed as previously described (*12*). In brief, the Cell Ranger software pipeline (version 5.0.1, 10x Genomics, Pleasanton, CA) was used to demultiplex cellular barcodes and map reads to the reference genome (refdata-cellranger-hg19-1.2.0 (human) or refdata-gex-mm10-2020-A (mouse). Seurat (version 4.0.2) demultiplexing function HTODemux was used to demultiplex the hash-tag samples. We filtered out the cells in which less than 500 genes or more than 6000 genes were detected. We further removed low-quality cells with more than 10% mitochondrial genes.

### Dimensionality reduction and clustering

The Seurat package (version 4.0.2) was used to perform unsupervised clustering analysis on scRNA-seq data (*43*). In brief, gene counts for cells were normalized by library size and log-transformed. To reduce batch effects, we apply the integration method implemented in the latest Seurat version 4 (function FindIntegrationAnchors and IntegrateData, dims = 1:30). The integrated matrix was then scaled by ScaleData function (default parameters). Principal component analysis was performed on the scaled data (function RunPCA, npcs = 30) in order to reduce dimensionality. 30 principal components were determined by using the ElbowPlot function and used to compute the KNN graph based on the euclidean distance (function FindNeighbors), which then generated cell clusters using function FindClusters. Uniform Manifold Approximation and Projection (UMAP) was used to visualize clustering results. The top differential expressed genes in each cluster were found using the FindAllMarkers function (min.pct = 0.1) and ran Wilcoxon rank sum tests. The differential expression between clusters were calculated by FindMarkers function (min.pct = 0.1), which also ran Wilcoxon rank sum tests.

### GM-CSF tissue score

A *GM-CSF tissue score* was calculated by selection of 25 genes related to GM-CSF-mediated immune response according to the literature published before: (*CSF2RA, CSF2RB, CXCL1, CCR2, CCR5, SHC1, GRB2, STAT5A, STAT5B, CXCR4, IL1B, IL6, TGFB1, FLT4, MAPK1, IRF4, IRF8, PIM1, SPI1, PPARG, CFH, CLEC5A, ALOX5AP, PPBP, CXCL5*). The *GM-CSF tissue score* was generated by transforming log2 expression profiles into Z-scores, and averaging Z-scores of the GM-CSF related gene sets to generate a tissue score for each patient sample.

### Statistics

Statistical analysis was performed using GraphPad Prism (La Jolla, CA). The results are shown as mean ± SEM when presented as a bar graph or as single data points with the mean in a scatter dot plot. Differences between two individual groups were compared using a two-tailed t test. In the case of three or more groups, a one-way ANOVA with Tukey’s multiple comparisons test was used. The correlation coefficient r was calculated using a Pearson correlation and the related P value was based on a t-distribution test.

## ACKNOWLEDGMENTS

FACS sorting was performed at the UKE FACS sorting core facility. Single-cell RNA sequencing was performed at the UKE Single-Cell Sequencing Core Facility. Next-generation sequencing was performed at the DFG-funded “Competence Centre for Genome Analysis Kiel” at the University of Kiel.

## CONTRIBUTIONS

Conceptualization: C.F.K. and U.P. Methodology: H.J.P., N.S., N.A., D.D.F., S.T., Y.Z., J. H.R., A.S., A.P., A.K. and A.B. Formal analysis: H.J.P., N.S., N.A., G.T., C.F.K, and U.P. Flow cytometry: H.J.P., N.A., S.T., D.D.F., scRNA sequencing: A.S., A.B., and C.F.K. Preclinical models: H.J.P., N.S., N.A., J.H.R., A.P., and A.K. Data analysis: H.J.P., N.S., Y.Z., M.H, Renal histology: N.A., A.P., C.M.-S., T.W., and U.P. Patient cohorts: U.O.W., J.H.R., O.M.S., E.M., M.T.L., E.H., R.A.K.S., T.B.H., T.W., C.F.K., and U.P. Writing original draft: C.F.K. and U.P. Writing review and editing: H.J.P., N.S., N.A., D.D.F., S.T., Y.Z., J.H.R, S.B., J.E.T., B.B., C.F.K., and U.P. Visualization: N.S., N.A., C.M.S., C.F.K., and U.P. Supervision: B.B., C.F.K. and U.P. Funding acquisition: T.B.H., S.B., C.M.S., B.B., C.F.K., U.P.

## DATA AVAILABILITY

The single-cell gene expression count and metadata tables containing clustering and quality control metrics for each cell will be publicly available at FigShare and via the Sequence Read Archive via GEO. All other data needed to evaluate the conclusions in the paper are contained in the paper or in Supplementary Materials.

