## Supplementary Figures for "GM-CSF drives immune-mediated glomerular disease by licensing monocyte-derived cells to produce MMP12"

| Cohort | Hamburg GN Registry Discovery I | European Renal cDNA Bank Discovery II | | | | CRU 228 ANCA cohort Validation | |
|---|---|---|---|---|---|---|---|
| **Disease** | **ANCA** | **Living donor** | **ANCA** | **Lupus nephritis** | **IgA nephropathy** | **Living donor** | **ANCA** |
| Patients (number) | 21 | 19 | 22 | 32 | 27 | 12 | 29 |
| Age (years) | 66.8 (±11.2) | NA | 58.69 (±14.13) | 35.06 (±13.26) | 36.40 (±14.07) | NA | 66.4 (±11.5) |
| Sex (% female / male) | 38.1 / 61.9 | NA | 45.5 / 54.5 | 78.1 / 21.9 | 29.6 / 70.4 | NA | 13.8 / 86.2 |
| Creatinine at time of biopsy (mg/dl) | 3.47 (±2.34) | NA | 2.43 (± 1.90) | 1.43 (±0.84) | 1.64 (±1.57) | NA | 3.07 (±2.76) |
| ANCA (cANCA / pANCA) | 9 / 12 | NA | 8 / 15 * | NA | NA | NA | 13 / 16 |
| Profiling (patient number) | FACS (9 / 5) [#]<br>scRNA-seq (8)<br>Pathology (19) | Transcriptome (19) | Transcriptome (22)<br>Pathology (19) | Transcriptome (32) | Transcriptome (27) | Transcriptome (12) | Transcriptome (29)<br>Pathology (29) |

NA: not announced; * one patient was tested positive for cANCA and pANCA ; [#] Chemokine receptor staining / intracellular cytokine staining

**Supplemental Table 1: Baseline characteristics of patients with ANCA-GN from different cohorts at the time of biopsy.**

Blood samples and renal biopsies were used for analyses in main Figure 1 and Figure 4. ANCA-GN = anti-neutrophil cytoplasmic antibody-associated glomerulonephritis; cANCA = cytoplasmic ANCA; pANCA= perinuclear ANCA.

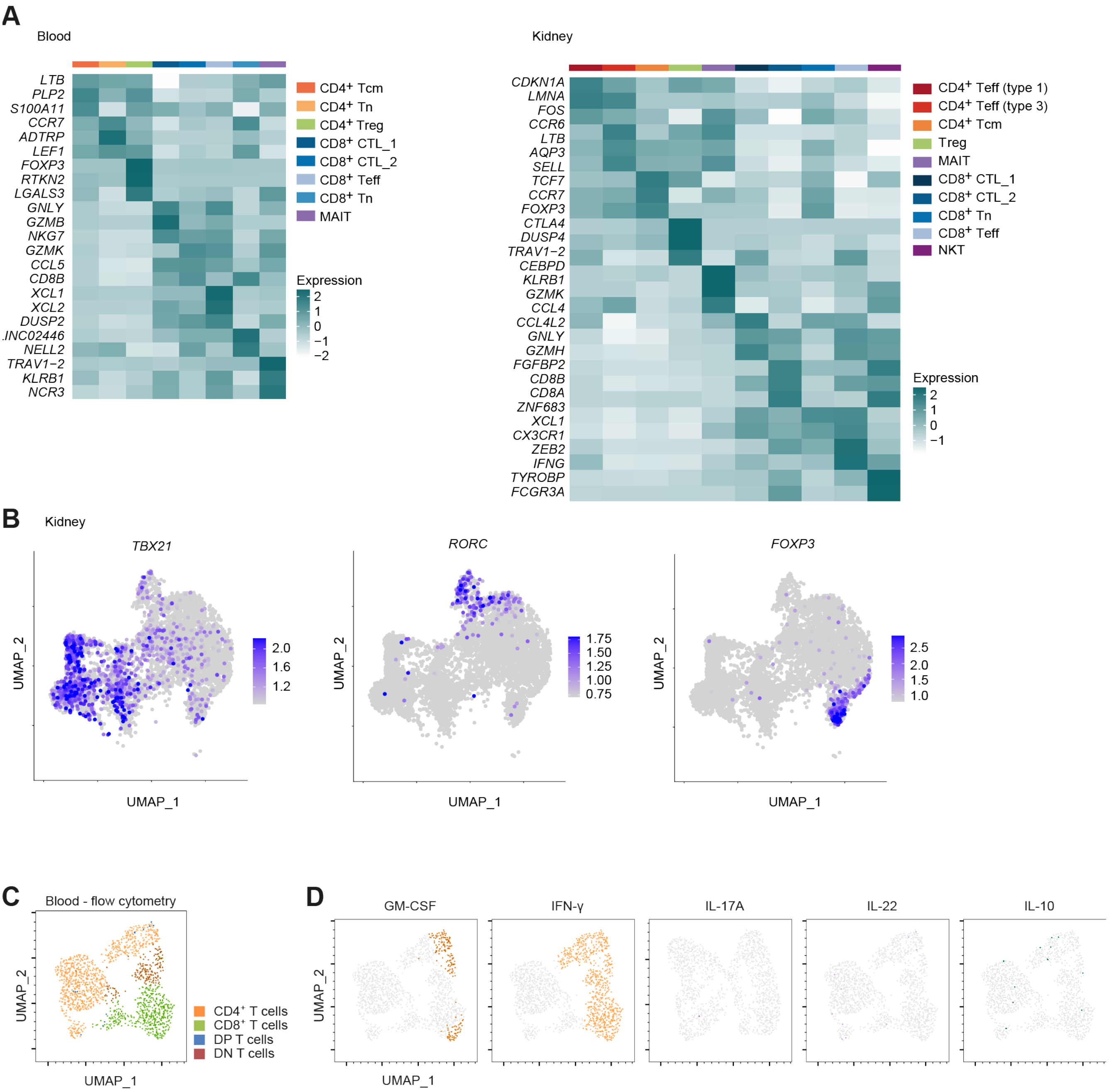


**Supplemental Figure 1: scRNA-seq analysis of T cells from blood and kidney tissue from ANCA GN patients.** **(A)** Human $CD3^{+}$ T cells were sorted from renal biopsies and blood samples and analyzed by scRNA-seq. Heatmaps show clustering of T cells according to their profile of differentially expressed genes. Clusters correspond to main Figure 1B. The heatmaps indicate the relative expression of individual genes in individual clusters ( blue-green: high expression; white: low expression). **(B)** UMAP plots showing the expression of indicated Th1 and Th17 marker genes in kidney (scale bars indicate normalized expression). **(C)** UMAP of FACS analyzed blood T cells and manually gated population identification as indicated (n=5). **(D)** Cytokine positive cells in the same UMAP.

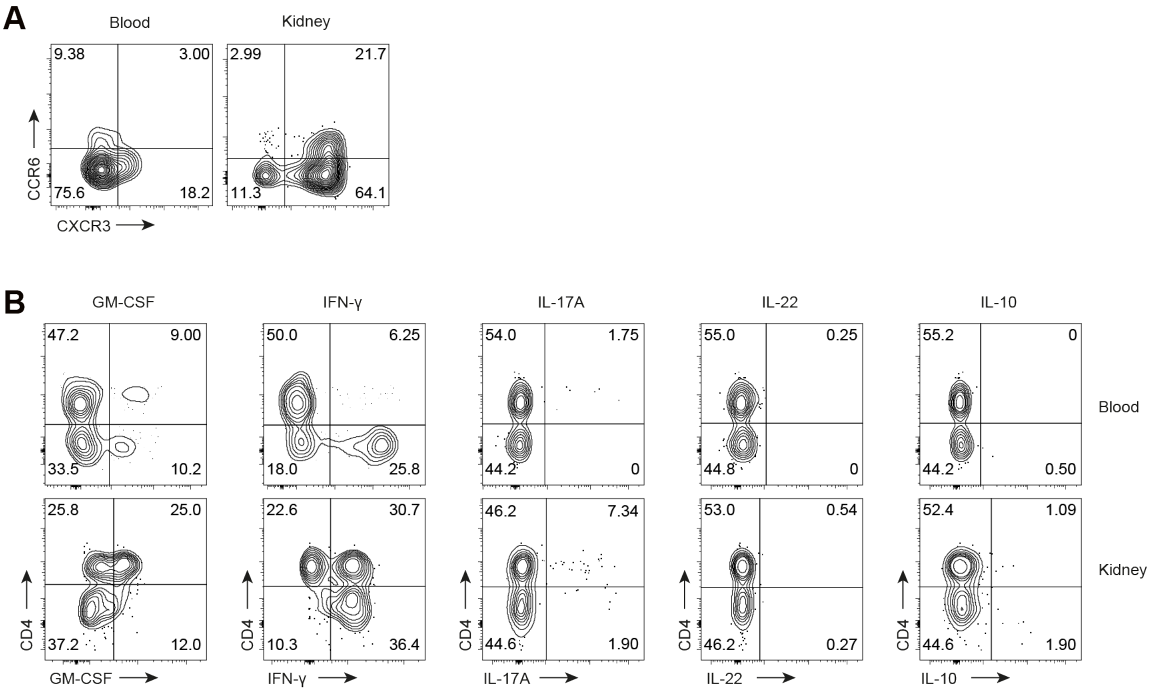


**Supplemental Figure 2: Identification and characterization of the type 1 immune response in the kidneys of patients with ANCA-GN.**

**(A)** Representative FACS data of CXCR3 and CCR6 expression in $CD3^{+}CD4^{+}$ T cells from renal biopsies and peripheral blood samples from ANCA-GN patients. **(B)** Representative FACS data of indicated cytokines produced in peripheral blood and kidney $CD3^{+}$ T cells of ANCA-GN patients after stimulation with PMA/ionomycin.

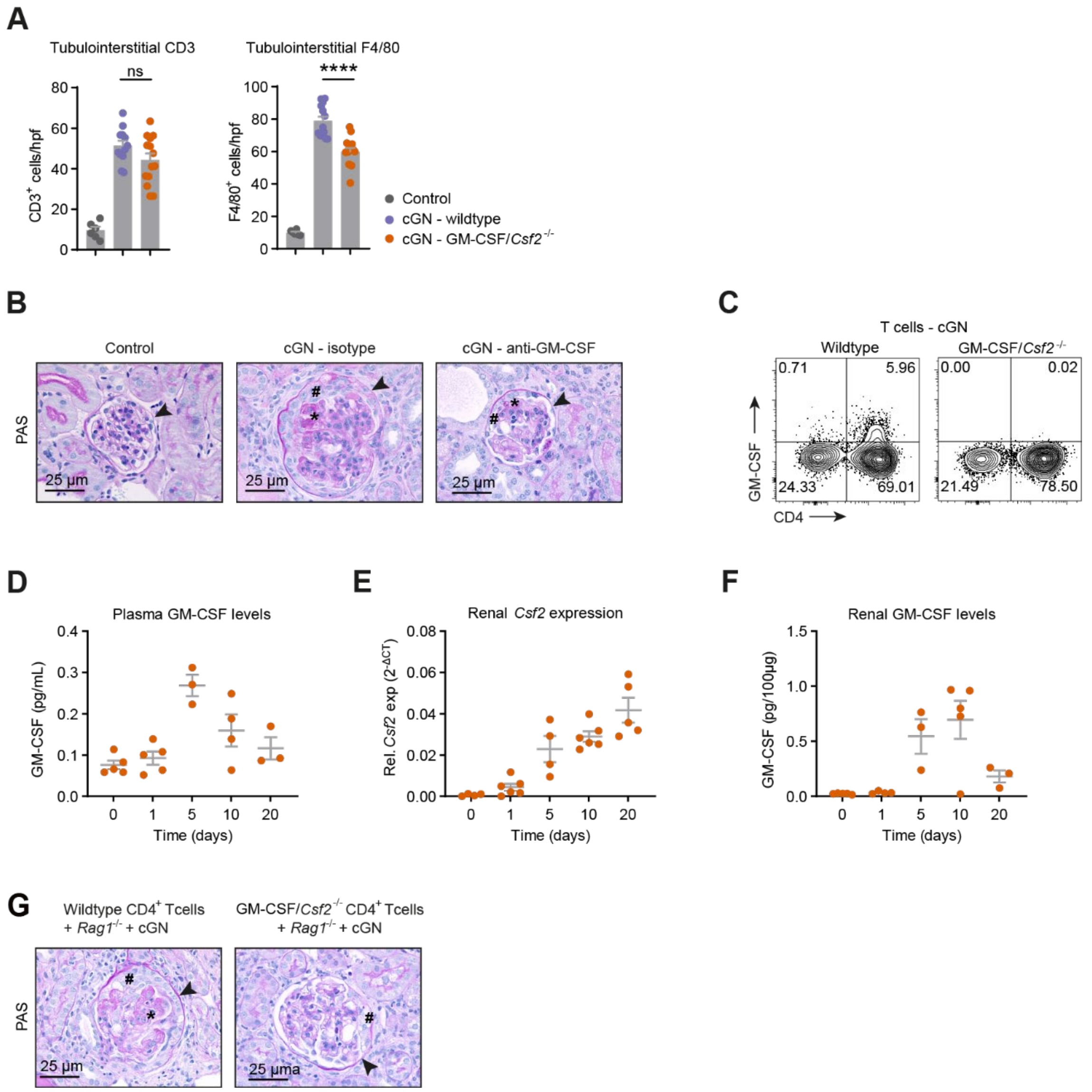


**Supplemental Figure 3: T cell-derived GM-CSF drives renal tissue damage in crescentic GN.**
**(A)** Quantification of tubulointerstitial CD3$^+$ and F4/80$^+$ cells per hpf (magnification x200) from wildtype and GM-CSF/*Csf2*$^{-/-}$ mice 10 days after cGN induction. **(B)** Representative photographs of PAS-stained kidney sections from control, nephritic wild-type mice treated with isotype antibody, or anti-GM-CSF antibody 10 days after cGN induction corresponding to main Figures 2D and 2E. **(C)** Flow cytometry analysis of stimulated renal lymphocytes from wildtype and GM-CSF/*Csf2*$^{-/-}$ mice 10 days after cGN induction. CD4$^+$T cells were assessed for GM-CSF production. **(D)** Quantification of GM-CSF protein in plasma from cGN mice at indicated timepoints. **(E)** Quantification of *Csf2* gene expression, as well as **(F)** GM-CSF protein in kidneys from cGN mice at indicated timepoints. **(G)** Representative photographs of PAS-stained kidney sections from Rag1$^{-/-}$ mice reconstituted with CD4$^+$ T cells isolated from spleens of wildtype or GM-CSF/*Csf2*$^{-/-}$ mice at day -1, analyzed 10 days after cGN induction corresponding to main Figures 2I and 2J.

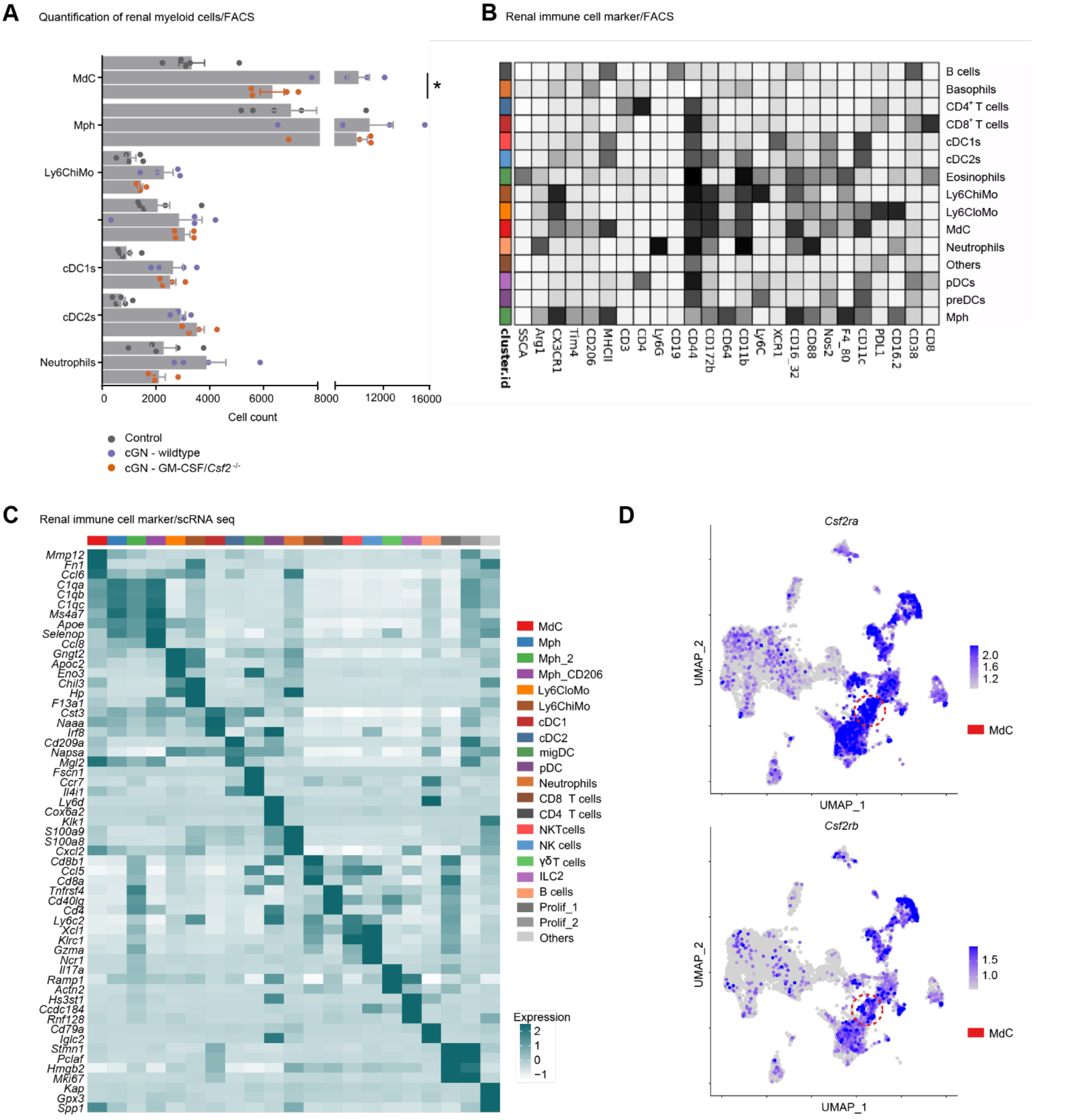


**Supplemental Figure 4: Profiling of renal immune cells after cGN induction.**
**(A)** Average number of renal myeloid cell populations from wildtype and GM-CSF/*Csf2*$^{-/-}$ mice 10 days after cGN induction. **(B)** Mean expression levels of all markers used for UMAP visualization and FlowSOM based clustering. Clusters correspond to main Figure 3A. **(C)** Mouse CD45$^{+}$ cells were sorted from kidneys and analyzed by scRNA-seq. Heatmaps show clustering of CD45$^{+}$ cells according to their profile of differentially expressed genes. Clusters correspond to main Figure 3B. **(D)** UMAP plots showing the expression of Csf2ra and Csf2rb in renal MdCs (scale bars indicate normalized expression).

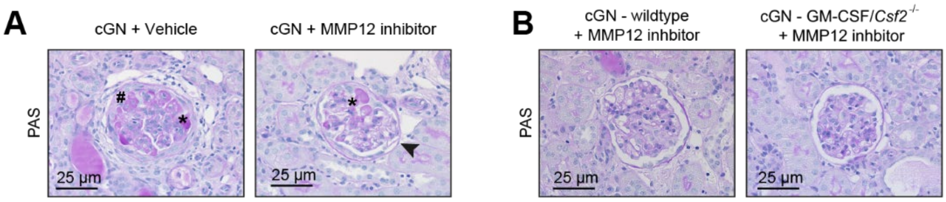


**Supplemental Figure 5: MMP12-driven kidney injury in cGN.**
**(A)** Representative photographs of PAS-stained kidney sections from nephritic wildtype mice treated with vehicle or pharmacological MMP12 inhibitor corresponding to main Figure 3J (► Bowman's capsule, * glomerular necrosis, # glomerular crescent). **(B)** Representative photographs of PAS-stained kidney sections from nephritic wildtype mice and GM-CSF/*Csf2*$^{-/-}$ mice treated with pharmacological MMP12 inhibitor corresponding to main Figure 3N.

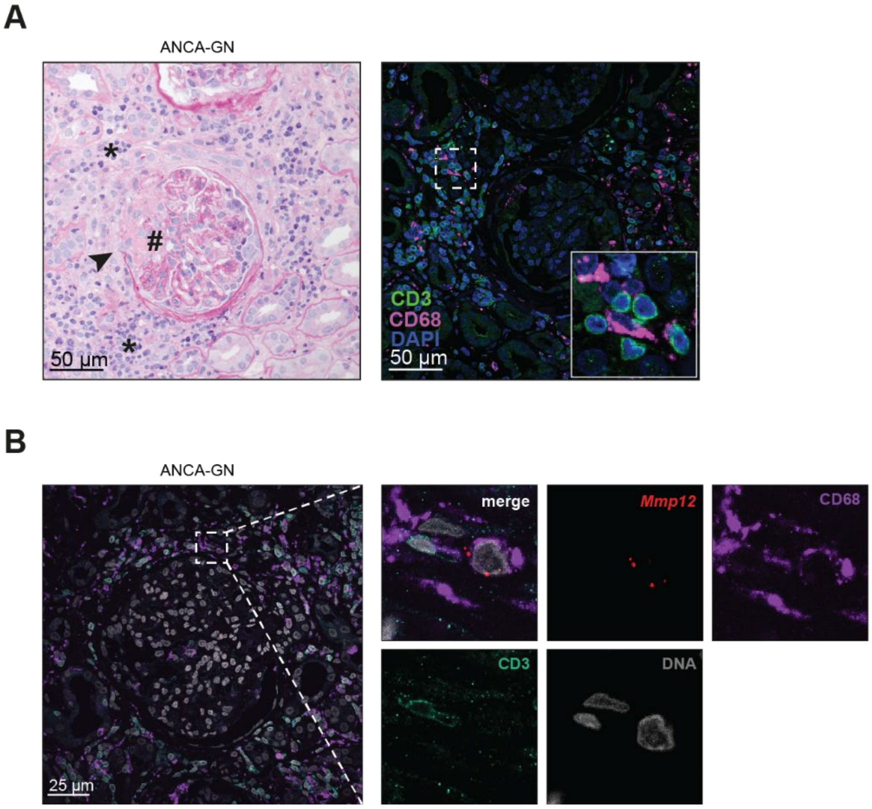


**Supplemental Figure 6: MMP12-expressing myeloid cells in renal biopsies from patients with ANCA-GN.**
**(A)** PAS and fluorescent staining for renal tissue histology of a patient with ANCA-associated glomerulonephritis. Left: rupture of the Bowman´s capsule (►), immune cell infiltration (*) and glomerular necrosis (#). Right: Immunofluorescence staining of the same glomerulus demonstrating the proximity of $CD3^+$ T cells and $CD68^+$ myeloid cells to the rupture. **(B)** Combined immunofluorescence staining of $CD3^+$ (green), and $CD68^+$ (violet) with FISH red (*MMP12*) of renal leukocytes in patients with ANCA-GN.

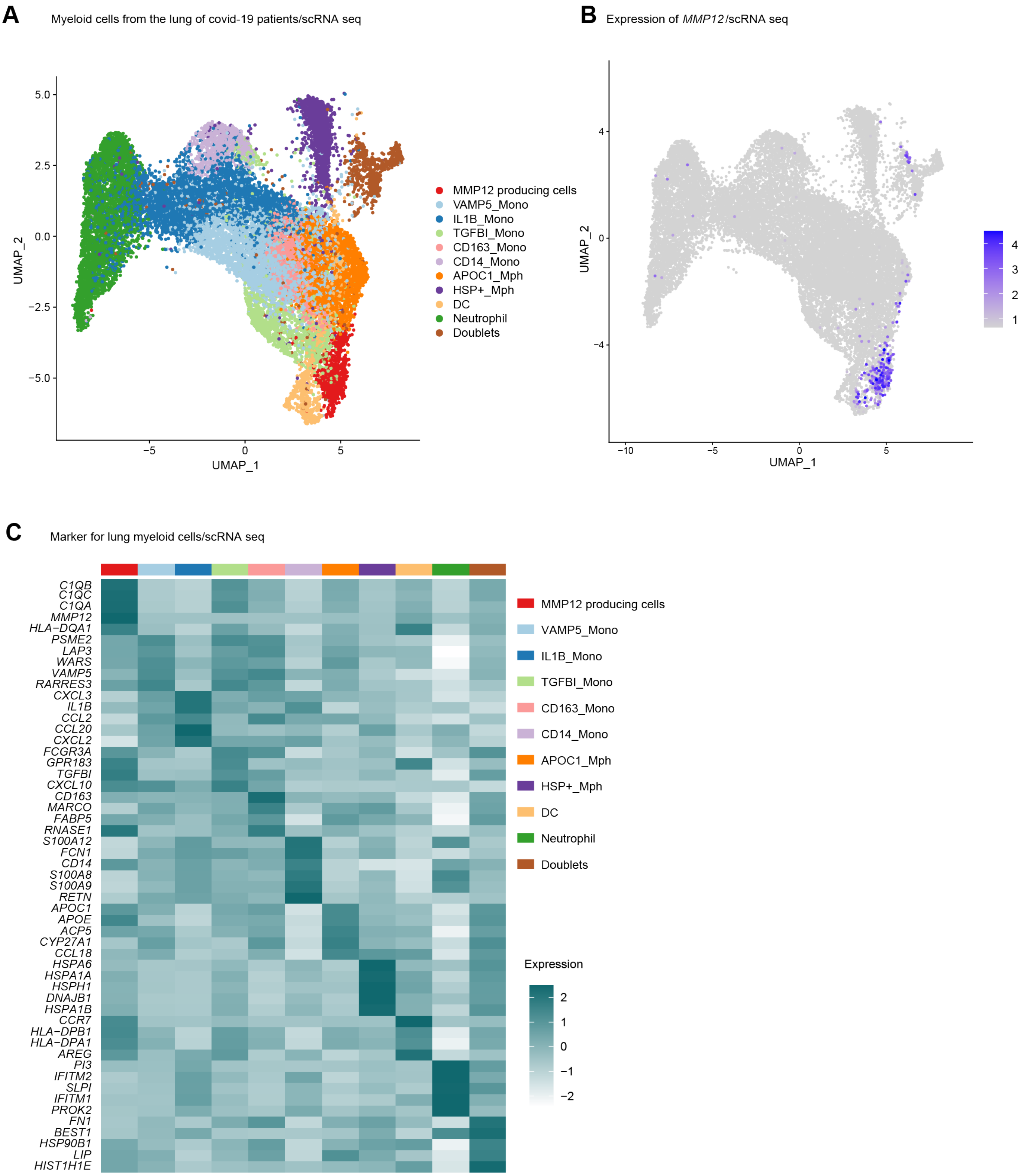


**Supplemental Figure 7: MMP12-expressing myeloid cells in the lungs of covid-19 patients.**
**(A)** UMAP dimensionality reduction embedding of myeloid cells from bronchoalveolar lavage fluid (BALF) of all patients from our study (COVID-19 n = 8). **(B)** UMAP plots showing the expression of *MMP12* (scale bars indicate normalized expression). **(C)** Heatmap showing clustering of myeloid cells according to their profile of differentially expressed genes. The heatmap indicates the relative expression of individual genes in individual clusters (blue-green: high expression; white: low expression).
